# BK current contributions to action potentials across the first postnatal week reflect age dependent changes in BK current kinetics in rat hippocampal neurons

**DOI:** 10.1101/839233

**Authors:** Michael Hunsberger, Michelle Mynlieff

## Abstract

The large conductance calcium-activated potassium (BK) channel is a critical regulator of neuronal action potential firing and follows two distinct trends in early postnatal development: an increase in total expression and a shift from the faster activating STREX isoform to the slower ZERO isoform. We analyzed the functional consequences of developmental trends in BK channel expression in hippocampal neurons isolated from neonatal rats aged one to seven days. Following overnight cultures, action potentials were recorded using whole-cell patch clamp electrophysiology. This population of neurons undergoes a steady increase in excitability during this time and the effect of blockade of BK channel activity with 100 nM iberiotoxin, changes as the neurons mature. BK currents contribute significantly more to single action potentials in neurons of one-day old rats (with BK blockade extending action potential duration by 0.46±0.12 ms) than in those of seven-day old rats (with BK blockade extending action potential duration by 0.17±0.05 ms). BK currents also contribute consistently to maintain firing rates in neurons of one-day old rats throughout extended action potential firing; BK blockade evenly depresses action potentials frequency across action potential trains. In neurons from seven-day old rats, BK blockade initially increases firing frequency and then progressively decreases frequency as firing continues, ultimately depressing neuronal firing rates to a greater extent than in the neurons from one day old animals. These results are consistent with a transition from low expression of a fast activating BK isoform (STREX) to high expression of a slower activating isoform (ZERO).

**New and Noteworthy:** This work describes the early developmental trends of BK channel activity. Early developmental trends in expression of BK channels, both total expression and relative isoform expression, have been previously reported, but little work describes the effect of these changes in expression patterns on excitability. Here, we show that early changes in BK channel expression patterns lead to changes in the role of BK channels in determining the action potential waveform and neuronal excitability.

## Introduction

The brain undergoes major developmental changes in excitability from infancy to adulthood. These include reorganization and establishment of circuits (Cline 2001; Gómez-Di Cesare et al. 1997; Huttenlocher and Dabholkar 1997), development of GABAergic and glutamatergic neurotransmitter systems (Dzhala and Staley 2003; Kumar et al. 2002; Szczurowska and Mareš 2013), and changes in the intrinsic excitability of cells resulting from establishment of mature patterns of voltage-gated ion channel expression (Gao and Ziskind-Conhaim 2017; Picken Bahrey and Moody 2003). Changes in intrinsic excitability are reflected in changes in the action potential waveform. Data from cortical pyramidal neurons, neocortical and cortical interneurons, and spinal motor neurons show that the neuronal action potential waveforms transition from longer lasting, low-amplitude spikes in immature neurons to high-amplitude, brief spikes accompanied by faster firing rates in mature neurons (Gao and Ziskind-Conhaim 2017; Goldberg et al. 2011; Kinnischtzke et al. 2012; McCormick and Prince 1987; Picken Bahrey and Moody 2003).

The large-conductance high-voltage and calcium-activated potassium channel (BK channel, Slo1, Maxi-K, Kcnma1) plays a central role in neuronal excitability and is also implicated in developmental changes in excitability. The BK channel is expressed along the axon and at presynaptic terminals, positioning it to regulate the frequency of action potential firing and neurotransmitter release (Jaffe et al. 2011; Misonou et al. 2006). BK channel expression patterns change through early development suggesting that its role in regulating excitability may be developmentally dependent. There is a several fold increase in total BK expression from late embryonic development to the first postnatal week in rodents; at the same time the predominant isoform shifts from STREX to the ZERO isoform (MacDonald et al. 2006). The STREX (STRess hormone-regulated EXon) isoform contains a 57 amino acid insert in the intracellular calcium sensing domain that confers higher calcium sensitivity and thus, requires lower calcium concentrations to shift the activation voltage to physiologically relevant potentials (Chen et al. 2005; Xie and McCobb 1998). Currents of the STREX isoform exhibit faster activation than those of the insertless ZERO isoform. In addition, the STREX insert changes the response of BK channels to phosphorylation so that phosphorylation by protein kinase A (PKA) causes inhibition of current in contrast to the typical increase in BK current seen in ZERO. In ZERO phosphorylation by protein kinase C (PKC) leads to a decrease in BK current whereas the STREX isoform is insensitive to PKC phosphorylation (Zhou et al. 2010, 2012).

While the developmental changes in expression have been described, investigation into the effects these changes have on neuronal properties and excitability is needed. The changes in BK channel expression and isoform characteristics suggest that the role of BK channels in modulating the action potential kinetics and firing rate changes as the nervous system matures. In the present study we used hippocampal neurons isolated from rats aged one to seven days to determine how excitability and the role of BK channels in this population of neurons change during this time as it is a period in which there are rapid changes in BK expression levels and isoforms.

We found, in agreement with earlier findings in cortical pyramidal neurons, cortical interneurons, and spinal neurons (Gao and Ziskind-Conhaim 2017; Goldberg et al. 2011; Kinnischtzke et al. 2012; McCormick and Prince 1987), that through early development hippocampal neurons become more excitable through early development as evidenced by greater firing rates and a transition from slow, low amplitude action potentials to fast, higher amplitude action potentials. Through this period, the effect of blocking BK channel activity changes. BK channel blockade has a greater effect on the duration of single action potentials at postnatal day one than at postnatal day seven despite large increases in total BK channel expression as development progresses. BK channel blockade also depresses firing rates to a greater extent at the onset of successive action potential firing in neurons from postnatal day one rats than in neurons postnatal day seven rats but has a greater effect in neurons from one-week-old rats as successive action potential firing continues.

## Materials and methods

### Laboratory animals

All animal protocols were approved by the Marquette University Institutional Animal Care and Use Committee according to the guidelines set forth by the National Research Council in the guide for care and use of laboratory animals. Sasco Sprague-Dawley rats (Charles River; Wilmington, MA) were housed and bred at Marquette University. Pups were removed from mothers just prior to tissue isolation.

### Isolation and culturing of hippocampal neurons

Acute primary cultures of hippocampal neurons were obtained as previously described (Mynlieff 1997). Briefly, the superior regions of the hippocampi were dissected from postnatal day 1 through 7 (P1-P7) rat pups in cold, oxygenated rodent Ringer’s solution (146 mM NaCl, 5 mM KCl, 2 mM CaCl_2_, 1 mM MgCl_2_, 10 mM HEPES, 11 mM D-glucose, adjusted to pH 7.4 with NaOH). Tissue was incubated in piperazine-N-N’-bis-2-ethanesulfonic acid (PIPES) buffered saline (120 mM NaCl, 5 mM KCl, 1 mM CaCl_2_, 1 mM MgCl_2_, 25 mM glucose, 20 mM PIPES, adjusted to pH 7.0 with NaOH) with 0.5% trypsin type XI (Sigma Aldrich, St. Louis, MO) and 0.01% DNase type I (Worthington, Lakewood, NJ) for 20-30 minutes. Enzymatic activity was terminated by rinsing the tissue with trypsin inhibitors (1 mg/mL trypsin inhibitor type II-O: chicken egg white and 1 mg/mL bovine serum albumin; Sigma-Aldrich, St. Louis, MO) in rodent Ringer’s solution. Following incubation, tissue was triturated to dissociate neurons and plated onto poly-L-lysine (30,000-70,000; Sigma-Aldrich, St. Louis, MO) coated culture dishes with Gibco neurobasal-A culture medium (ThermoFisher Scientific, Waltham, MA) fortified with B-27 supplement, 0.5 mM glutamine and 0.02 mg/mL gentamicin. Neurons were held in a 37 °C, 5% CO_2_ incubator overnight to allow for reinsertion of membrane proteins while minimizing *in vitro* developmental changes.

### Electrophysiology

Recordings were taken from cultured neurons the day after isolation to minimize the outgrowth of processes. Action potentials were obtained by whole-cell patch-clamp recording in current-clamp mode with a Dagan 3900A patch clamp amplifier (Dagan Corporation, Minneapolis, MN), Axon Digidata 1322A 16-bit data acquisition system, and pClamp 10.4 data acquisition software (Molecular Devices, San Jose, Ca). Thick walled borosilicate pipettes (≈4-8 MΩ) were filled with internal solution (140 mM K-Gluconate, 0.5 mM CaCl_2_, 2 mM MgCl_2_ 1 mM EGTA, 2 mM ATP-Na_2_, 0.2 mM GTP-Na_2_, and 10 mM HEPES, adjusted to pH of 7.2-7.4 with KOH). Cells were placed in an extracellular recording solution (115 mM NaCl, 25 mM NaHCO_3_, 2.5 mM KCl, 2 mM CaCl_2_, 1 mM MgCl_2_, 10 mM glucose, 10 mM HEPES, adjusted to pH of 7.4 with NaOH). Single action potential were elicited by a 0.1 ms, 8 nA current injection and action potential trains were elicited by 100 ms and 1 s depolarizing current injections ranging from 10-200 pA until a maximum number of action potentials was evoked. Data was digitized at a rate of 20 kHz with a 10 kHz low pass filter. All membrane potentials reported have been adjusted post-recording to account for the liquid junction potential (15.4 mV).

To assess the contribution of BK channels to action potentials 100 nM Iberiotoxin (Alomone Labs, Jerusalem, Israel) was administered using a U-tube drug delivery system consisting of PE-10 polyethylene tubing with a single perforation to allow transient, focal drug application.

### Drug Preparation

Iberiotoxin (IbTx; Alomone Labs, Jerusalem, Israel) was dissolved to a 1 mM intermediate stock in extracellular recording solution and stored at −20 °C in multiple aliquots to minimize freeze-thaw cycles. 1 mM IbTx stock was thawed and added to the extracellular recording solution the same day as it was used for recording to prepare a 100 nM working solution (*K*_*i*_ = 250 pM-1 nM; (Galvez et al. 1990; Koschak et al. 1997).

### Data Analysis

Action potential durations were calculated as the time interval between the rising and falling phase at one-half of the spike amplitude. Fast afterhyperpolarization (fAHP) magnitude was measured as the lowest voltage attained during hyperpolarization subtracted from the resting membrane potential recorded prior to the stimulus used to induce the action potential. For the IbTx-induced differences in action potential duration, fAHP, and the maximum number of action potential evoked in 100 ms the reported values were calculated as the difference between the value measured from the IbTx-treated trace and the average value measured from one untreated trace before, and one after IbTx application to account for potential confounding effects of prolonged patching and electrical stimulation. Instantaneous frequencies were calculated as the inverse of the peak-to-peak time interval between two successive action potentials.

All statistical tests were performed using SigmaPlot 14 (Systat Software, San Jose, Ca). Much of the data collected was not normally distributed, therefore non-parametric statistical tests were used where appropriate. Data were initially separated by sex, but ultimately pooled as no statistical differences by sex were observed.

## Results

Data were collected from dissociated heterogeneous cultures of neurons from the superior hippocampi of rats aged one to seven days. Unlike cultures from embryonic rodents which are enriched with pyramidal neurons these cultures have a large percentage of interneurons (Aika et al. 1994; Banker and Cowan 1979; Mynlieff 1997, 1999). For all recordings, current was injected to adjust membrane potentials to −65 to −70 mV prior to collecting action potential measurements to normalize the electrochemical driving force for all cells. We found that the mean action potential amplitude increased while the duration decreased as the neurons matured (**Fig 1**). The mean action potential amplitude increased from 80.96±1.48 mV (n=80 cells, 9 animals) in neurons isolated from one day old (P1) rats to 91.65±1.39 mV in neurons isolated from seven day old (P7) rats (n= 71 cells, 11 animals; one-way ANOVA on Ranks, p<0.001, Dunn’s Multiple comparisons, p<0.05; **Fig 1B**). While the amplitude significantly increased, the amplitude remained highly variable, reflecting a diverse cell population; amplitudes ranged from 53.65 to 107.76 mV in P1 neurons to 57.71 to 108.81 mV in P7 neurons (**Fig 1C**).

**Figure 1:**
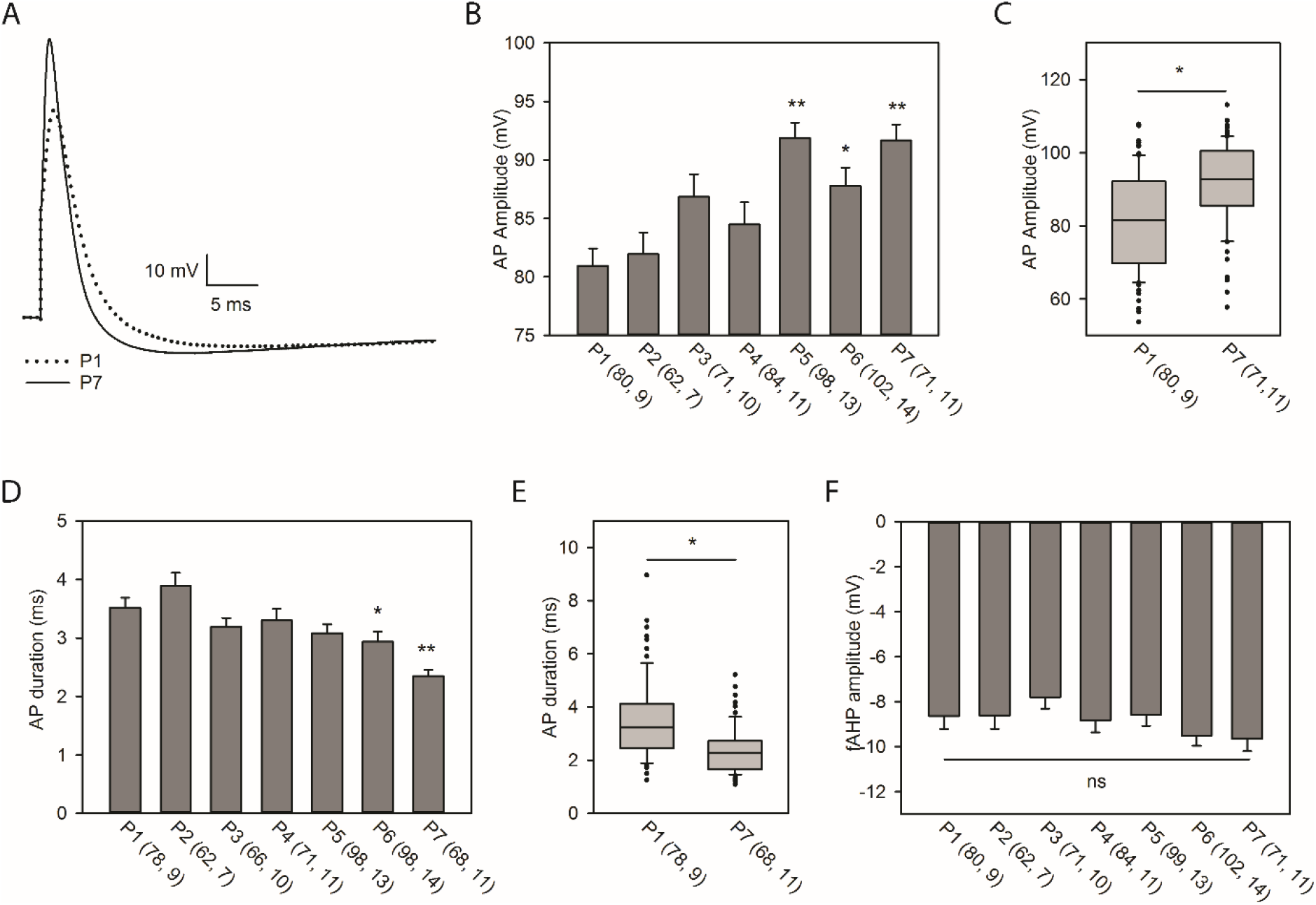
The action potential waveform changes across the first postnatal week. **(A)** Representative traces comparing a typical action potential from a P1 neuron (dotted trace) to that of a P7 neuron (solid trace). **(B)** Mean+SEM of action potential amplitudes for P1 to P7 neurons. A single asterisk indicates significant difference from P1; a double asterisk indicates significant difference from P1 and P2 (one-way ANOVA on ranks, p<0.001, Dunn’s Multiple comparisons, p<0.05). **(C)** Box and whisker plot of P1 and P7 action potential amplitudes is depicted to display the variability of the data. The box represents the second-third quartiles with the median indicated by the line; the whiskers represent the 10^th^-90^th^ percentile. **(D)** Mean+SEM action potential duration for P1 to P7 neurons measured at half-maximal amplitude. A single asterisk indicates significant difference from P1 and P2; a double asterisk indicates significant difference from P1-P5. (one-way ANOVA on ranks, p<0.001, Dunn’s multiple comparisons p<0.01). **(E)** Box and whisker plot of P1 and P7 action potential durations is depicted to display the variability of the data. The box represents the second-third quartiles with the median indicated by the line; the whiskers represent the 10^th^-90^th^ percentile. **(F)** Mean+SEM of fast afterhyperpolarizations (fAHP). For B, D, and F, the first number in parentheses is the number of individual neurons and the second number is the number of rats from which the neurons were obtained.

During the same period the mean duration, measured as the width of the action potential at half of the amplitude, decreased from 3.52±0.17 ms (P1, n=78 cells, 9 animals) to 2.35±0.11 ms (P7, n=68 cells, 11 animals; one-way ANOVA on ranks, p<0.001, Dunn’s multiple comparisons p<0.01; **Fig 1D**). As in the case of action potential amplitude, action potential durations were highly variable, ranging from 1.25 to 8.95 ms in P1 neurons and from 1.08 to 5.21 ms in P7 neurons (**Fig 1E**).

This overall sharpening of the action potential waveform likely reflects an overall increase in ion channel expression, with an increase in voltage gated Na^+^ channels resulting in a faster depolarization and greater amplitude and an increase in voltage gated K^+^ channels resulting in faster repolarization (Gao and Ziskind-Conhaim 2017; Picken Bahrey and Moody 2003). In line with reports from other cell types (Goldberg et al. 2011; Kinnischtzke et al. 2012), the mean membrane resistance of these neurons decreased from 1.93±0.13 GΩ in P1 neurons (n=80 cells, 9 animals) to 1.03±0.11 GΩ in P7 neurons (n=70 cells, 11 animals; t-test, P<0.001) reflecting an increase in ion channel expression with no change in the cell capacitance, which reflects cell size (**Table 1**). No significant changes were observed in the magnitude of the fast afterhyperpolarization (fAHP; **Fig 1F**). As the neuronal population from which recordings were obtained is highly diverse, it is possible that any age-dependent changes were obscured by the variability of the data due to cell types.

**Table 1:** Membrane properties of P1 and P7 cultured hippocampal neurons. Mean±SEM of the resting membrane potential, membrane resistance, and the capacitance of cultured P1 and P7 superior hippocampal neurons. Membrane resistances have been adjusted from the raw data to reflect the liquid junction potential. Statistical comparisons were made using a two sample t-tests.

|  | P1 | P7 | t-test |
| --- | --- | --- | --- |
| **Resting membrane potential (mV)** | -57.9±1.3 (n=72) | -56.9±1.0 (n=70) | ns |
| **Membrane Resistance (GΩ)** | 1.93±0.13 (n=80) | 1.03±0.11 (n=70) | p<0.001* |
| **Capacitance (pF)** | 21.2±0.7 (n=72) | 21.9±0.7 (n=70) | ns |

Changes in the waveform of action potentials can change the excitability of the cell as a shorter duration action potential can enable a higher firing frequency (Picken Bahrey and Moody 2003). Therefore we compared the excitability, represented by the maximal number of action potentials that could be evoked by a 100 ms depolarizing pulse, in neurons from heterogeneous hippocampal cultures across the first postnatal week. A range of pulses from 10-200 pA were injected into the cells and the maximum number of action potential elicited by a single pulse is reported. P1 neurons fired a maximum of 2.0±0.1 action potentials (n=80 cells, 9 animals), while P7 neurons fired 3.3±0.2 action potentials (n=71 cells, 11 animals; one-way ANOVA on ranks, p<0.001, Dunn’s multiple comparisons, p<0.05; **Fig 2**) The variability of P1 and P7 is shown in **Fig 2C**.

**Figure 2:**
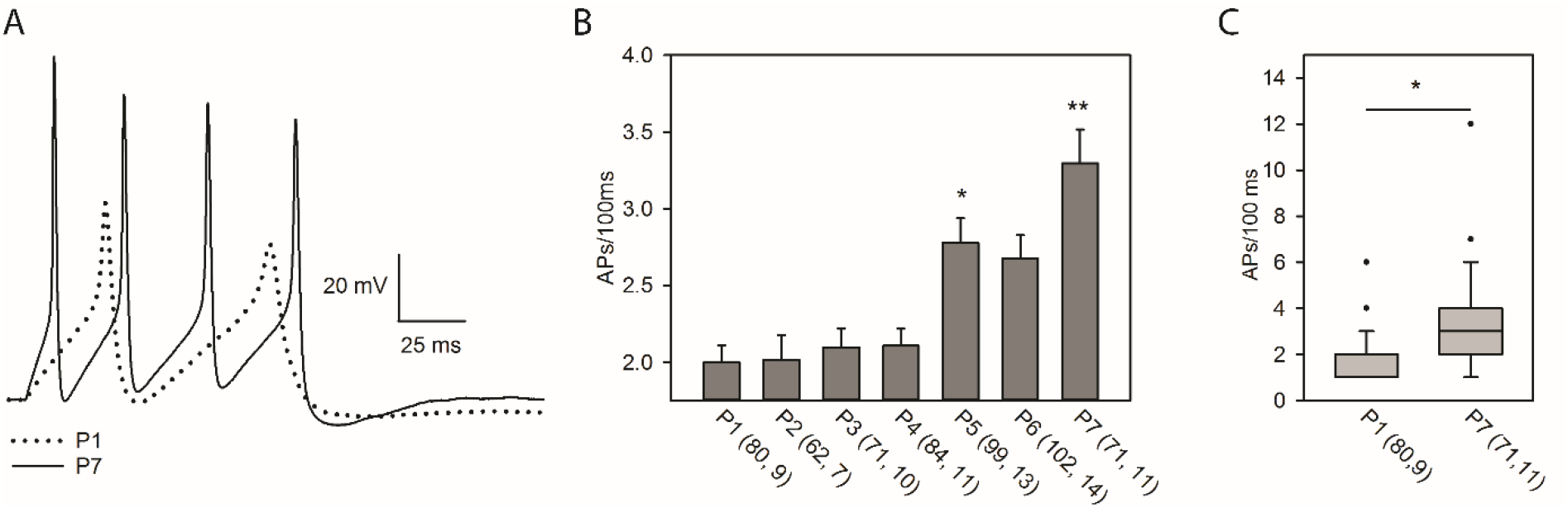
Excitability of hippocampal neurons increases across the first postnatal week. **(A)** Representative traces comparing action potential firing in response to a 100 ms depolarizing pulse in a P1 (dotted) and P7 neuron (solid). **(B)** Mean+SEM of maximal action potential firing per 100 ms for P1 to P7 neurons. A single asterisk indicates significant difference from P1-2; a double asterisk indicates significant difference from P1-4 (one-way ANOVA on ranks, p<0.001, Dunn’s multiple comparisons, p<0.05). **(C)** Box and whisker plot of P1 and P7 maximal action potentials fired per 100 ms. The box represents the second-third quartiles with the median indicated by the line; the whiskers represent the 10^th^-90^th^ percentile.

To determine how BK channels contribute to action potentials across early development, we measured the effect of the selective BK channel antagonist iberiotoxin (IbTx, 100 nM) in P1-P7 neurons. The effect on duration was calculated as the difference between the action potential half-width during IbTx perfusion and the average of two control traces, one before perfusion and one after, to account for cytoplasmic washout from the recording pipette. Throughout the first postnatal week there was less effect of IbTx on action potential duration as the neurons matured (**Fig 3B**). In the presence of IbTx the duration of action potentials of P1 neurons increased by 0.63±0.10 ms (n=64 cells, 6 animals), while the duration of action potentials in P7 neurons only increased by 0.24±0.08 ms (n= 45 cells, 6 animals; one-way ANOVA, p=0.001, Dunn’s multiple comparisons, p<0.05). It is possible that the decrease in the effect of IbTx is the result of the decrease of the action potential duration with postnatal age rather than a result of different activity of BK channels at these two time points; the longer an action potential lasts, the longer the membrane is sufficiently depolarized for BK channels to open and IbTx will have a greater effect simply because more BK channels have opened. To investigate this possibility, we compared the effect of IbTx on action potential duration only in neurons with an initial (untreated with IbTx) action potential half-width of 2-3 ms. In this subset of P1-P2 neurons IbTx perfusion increased mean action potential duration by 0.46±0.12 ms (n=27 cells, 12 animals) while it only increased the mean action potential durations in P6-P7 neurons by 0.17±0.05 ms (n=47 cells, 12 animals; p=0.035; Mann-Whitney rank sum test; **Fig 3C**). These data indicate that the age-dependent change in the effect of IbTx on action potential duration in hippocampal neurons is the result of differences in BK channel activity across the first postnatal week and not merely a consequence of changes in the overall action potential kinetics during this time.

**Figure 3:**
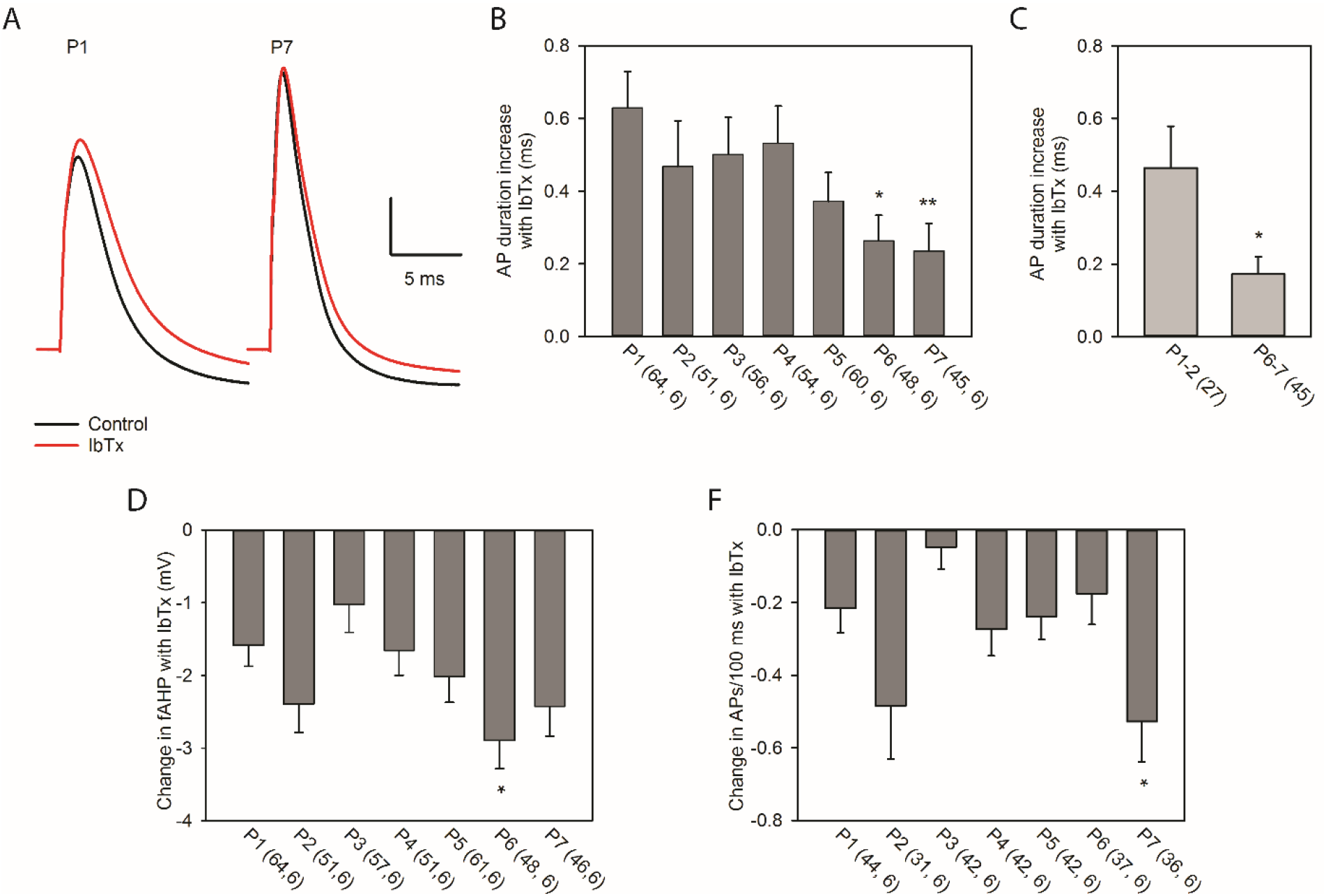
Iberiotoxin increases action potential duration to a greater extent in P1 neurons than in P7 neurons. **(A)** Representative traces comparing the effect of IbTx perfusion (control is in black, IbTx treatment is in red) on action potentials of P1 and P7 neurons. **(B)** Mean+SEM effect of IbTx on action potential duration. A single asterisk indicates significant difference from P1; a double asterisk indicates significant difference from P1 and P4. (one-way ANOVA on ranks, p=0.001, Dunn’s multiple comparisons, p<0.05). **(C)** Mean+SEM effect of IbTx on action potential duration in P1-P2 and P6-P7 neurons with an initial action potential duration of 2-3 ms. (Mann-Whitney rank sum test, p<0.05) **(D)** Mean+SEM magnitude of the effect of IbTx on the fast afterhyperpolarization (fAHP) following a single action potential. A single asterisk indicates significant difference from P3 (one-way ANOVA, p<0.01, Holm-Sidak multiple comparisons, p<0.01). **(E)** Mean+SEM effect of IbTx on the number of action potentials fired in 100 ms for all neurons capable of firing >1 action potentials. Asterisk indicates significant difference from P3 (one-way ANOVA on ranks, p<0.01, Dunn’s multiple comparisons, p<0.05). For B, D, and E, the first number in parentheses is the number of individual neurons and the second number is the number of rats from which the neurons were obtained.

We next examined the effect of IbTx on the fAHP (**Fig 3D**). While IbTx reduced the magnitude of the fAHP to a greater extent in P6 than in P3 neurons, it is unclear whether this is part of a broader trend, as no significant differences were observed between any other pairs of time points. Because these recordings were of randomly selected cells in heterogeneous cultures, it is possible that developmental trends were obscured by the variability of the data. To address this, we analyzed the effect of IbTx on fAHP magnitude within groups of cells with similar electrophysiological properties. Action potential data was binned both by initial action potential duration (1-2, 2-3, and 3-4 ms) and by initial magnitude of the fAHP (0-5, 5-10, and 10 to 15 mV). No age dependent changes within these groupings were observed (data not shown).

An important physiological role of BK channels is to enable high frequency firing (Gu et al. 2007). To examine how the role of BK channels in maintaining action potential firing frequency changes in postnatal development, fast firing hippocampal neurons (initial firing rate >20HZ with sustained firing throughout the 1s depolarization) were perfused with 100 nM IbTx while action potential trains (**Fig 4A)** were evoked by a series of ten one second depolarizing pulses from 10-100 pA. P1 neurons reached maximal firing rates at lower current inputs when compared to P7 neurons. IbTx had a significant effect on the input-output relationship (2-way ANOVA, p<0.001 for the main effect of IbTx for both ages tested; **Fig 4B**).

**Figure 4:**
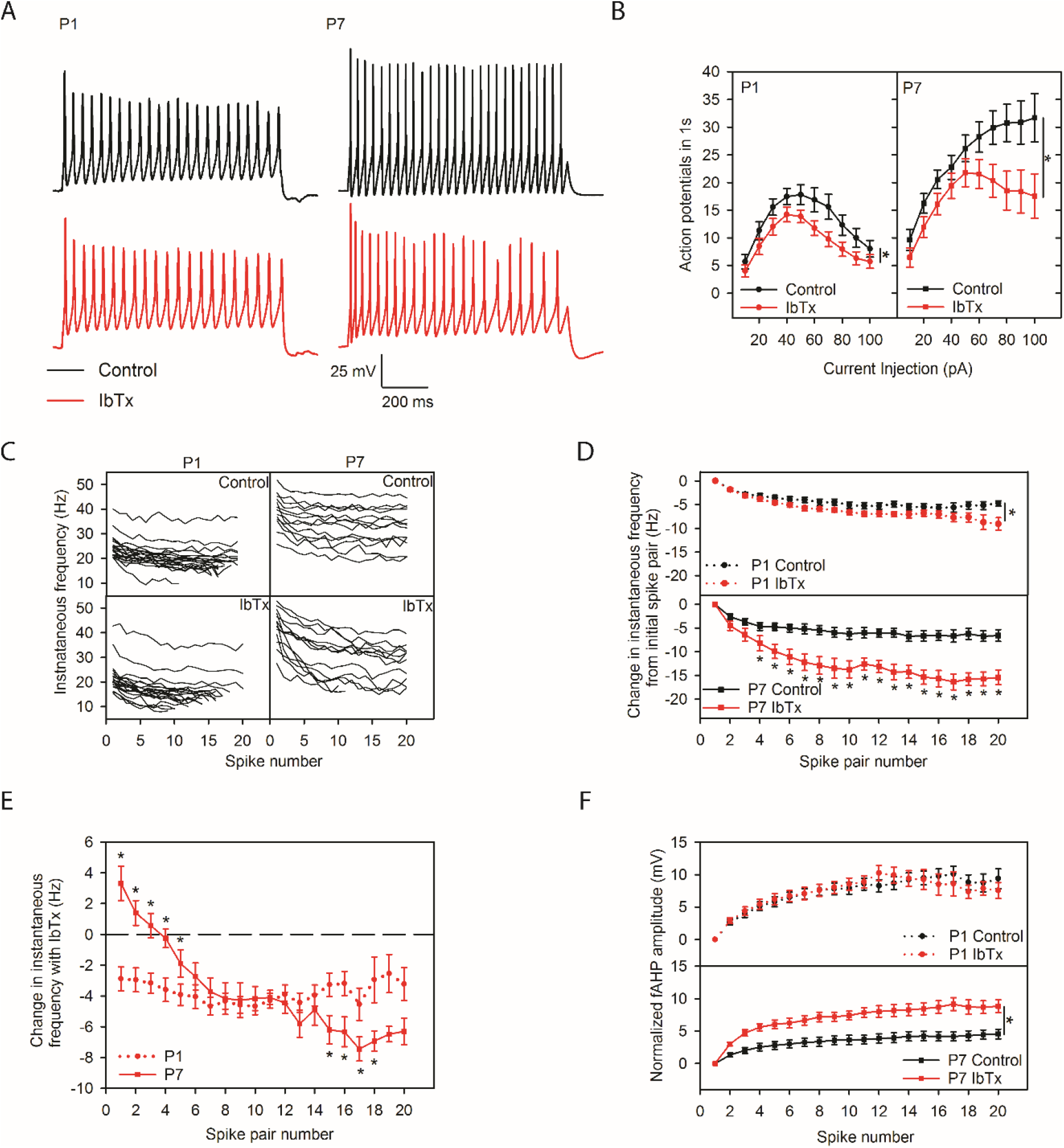
Iberiotoxin effects have different timing in P1 and P7 neurons. **(A)** Representative traces of 1s action potential trains from P1 and P7 neurons before (black) and after IbTx perfusion (red). **(B)** Input-output curves for P1 (left) and P7 (right) without (black trace) and with IbTx (red trace) for 10-100 pA current injections. Asterisks at the end of the curves indicate a main effect of IbTx (two-way ANOVA, p<0.001 for the main effect of IbTx). **(C)** Spike-frequency plots over the first twenty spike pairs for all cells included in the analysis of fast-firing neurons. **(D)** Mean±SEM change in f_i_ from the initial f_i_ for the first twenty spike pairs for P1 and P7 fast firing hippocampal neurons before and after IbTx application. The asterisk at the end of the P1 curves indicates that there is a main effect of IbTx on successive firing frequency (two-way ANOVA, p<0.001 for the main effect of IbTx). Asterisks comparing individual data points in the P7 curves indicate a significant interaction of IbTx and the spike pair number (two-way ANOVA, p=0.002, Holm-Sidak pairwise comparison, p<0.05). **(E)** Mean±SEM effect of IbTx (the difference between f_i_-IbTx and f_i_-control) on the f_i_ of the first 20 spike pairs. Asterisks indicate significant differences between the effect of IbTx on P1 and P7 neurons at that spike pair (two-way ANOVA, p<0.001, Holm-Sidak multiple comparisons, p<0.05). **(F)** Mean±SEM of the fAHP amplitude following the first 20 action potentials normalized to the amplitude of the first fAHP. The asterisk at the end of the P7 curves indicates a main effect of IbTx on fAHP amplitude (two-way ANOVA, p<0.001 for the main effect of IbTx).

We analyzed the maximally firing action potential train for each cell. We measured how instantaneous firing frequency (f_i_) decreased through the first twenty pairs of action potentials, measured as the difference in f_i_ from the initial f_i_ for each spike pair, for trains of action potentials from P1 and P7 neurons before and after IbTx application. Spike frequency plots of the first twenty action potential pairs for all neurons included are illustrated in **Fig 4C.** Neurons from P1 and P7 rats with and without IbTx showed some degree of spike-frequency adaptation; f_i_ decreased progressively throughout the action potential train for all conditions (**Fig 4D**). P1 neurons displayed similar spike-frequency adaptation in the presence and absence of iberiotoxin (**Fig 4D**, **top panel**) with the effect of IbTx having a modest effect on instantaneous frequency (two-way ANOVA, p<0.001 for the main effect of IbTx) although no significant differences could be detected for individual spike pairs. In contrast, IbTx perfusion of P7 neurons decreased f_i_ to a greater degree as action potential firing progressed (interaction of IbTx and spike pair number: p<0.001, two-way ANOVA, Holm-Sidak multiple comparisons, p<0.05), demonstrating that BK channels play a significant role in inhibiting spike frequency adaptation in more mature neurons (**Fig 4D**, **bottom panel**).

IbTx-induced changes in f_i_ within the first twenty action potential pairs were analyzed. In P1 neurons (n=20 cells, 9 animals) IbTx decreased action potential firing frequency throughout the action potential train compared to control recordings without IbTx. In P7 neurons (n=13 cells, 4 animals) IbTx significantly increased the f_i_ of the first spike pair and decreased f_i_ to an increasingly greater extent as the train progressed. IbTx reduced f_i_ to a greater extent in P1 neurons than in P7 neurons for the first five action potential pairs, while reducing f_i_ to a greater extent in cells from P7 animals than in cells from P1 animals for action potential pairs 15-18 (two-way ANOVA, p<0.001, Holm-Sidak multiple comparisons, p<0.05; **Fig 4E**). Not all cells fired enough action potentials to allow analysis of twenty spike pairs, so statistical power was reduced for later spike pairs. We also tested IbTx had a different effect on the afterhyperpolarization following each action potential (**Fig 4F**). IbTx perfusion caused afterhyperpolarizations to reach less negative membrane potentials in P7 fast firing neurons (**Fig 4F**, **bottom panel**), but not in P1 fast firing neurons (**Fig 4F**, **bottom panel**).

Among cells that met the defined criteria for fast firing cells, the P7 neurons fired significantly faster (**Fig 4C)**; the initial f_i_ for P1 neurons was 25.13±1.05 Hz versus 39.54±1.92 Hz for P7 neurons(p<0.01). To test whether the conclusions drawn from these experiments were appropriate, the analyses were repeated with a smaller subset of the data: the 6 fastest firing P1 neurons and the 6 slowest firing P7 neurons. Their initial mean f_i_’s were 30.38±2.20 Hz and 33.79±1.89 Hz (p=0.27), respectively and results of the full data set were preserved in this subset (data not shown), supporting the conclusion that the differences observed were age-dependent.

## Discussion

We found that excitability of hippocampal neurons increases through the first postnatal week, with the action potential waveform increasing in amplitude and decreasing in duration (**Fig 1**) while neurons become capable of firing more successive action potentials before entering depolarizing block (**Fig 2**). Our data also demonstrate that as development progresses the contribution of BK channels to the repolarization of single action potentials decreases (**Fig 3**). At the same time there is a shift in the timing of BK channel currents in high frequency firing, such that BK currents contribute to spike-frequency adaptation and afterhyperpolarizations in P7, but not P1 neurons (**Fig 4**).

Our findings on the increases in intrinsic excitability and changes in the action potential waveform through early development are consistent with previous reports in other neuronal populations (Gao and Ziskind-Conhaim 2017; Goldberg et al. 2011; Kinnischtzke et al. 2012; McCormick and Prince 1987). Prior data on changes in BK channel activity in this time period are limited, but one report found that channel density increases from postnatal day one to day twenty-eight and expression studies show that BK mRNA expression increases sharply throughout the central nervous system during the first postnatal week (Kang et al. 1996; MacDonald et al. 2006). While it may appear contradictory to observe greater contributions of BK channels to single action potentials in early development, this is sufficiently explained by a shift from fast activating BK channel variants to slow activating variants. Macdonald et. al. report a shift in the dominant hippocampal BK isoform from STREX, a more Ca^2+^-sensitive, faster acting variant than the slower ZERO isoform, a variant that may have too long of an activation time to influence the waveform of a single action potential (Chen et al. 2005; MacDonald et al. 2006; Xie and McCobb 1998). This interpretation is also supported by the change in timing of IbTx effects in prolonged firing observed in this report.

Beta subunit association can also affect the BK channel’s activation timing. The β-4 subunit, a neuronally expressed β subunit, increases the Ca^2+^ sensitivity but also slows the kinetics of the BK channel such that it would not participate in repolarization of a single action potential (Behrens et al. 2000). Expression of β-4 subunit mRNA (Kcnmb4) increases in the hippocampus with postnatal development making changes in beta subunit association another avenue to explore in examining postnatal changes in BK channels activity. However, it is unlikely that this is the source of the changes that we observed as we used the BK antagonist IbTx to examine BK channel activity; BK channels associated with the β-4 subunit are reportedly insensitive to blockade with IbTx as well as charybdotoxin (Meera et al. 2000).

The early postnatal period is notable for being a period of heightened seizure susceptibility (Rakhade and Jensen 2009). This is likely due to many reasons as the brain undergoes extensive developmental changes after birth. These reasons may include immature patterns of BK channel expression as described here. This is supported by findings that BK channel dysfunction can be causative in epilepsy. Administration of the BK channel antagonist paxilline reduces the incidence and duration of seizures in immature mice (Sheehan et al. 2009). One mutation in the intracellular RCK domain of the BK channel, D343G, has been identified in clinical settings as causative in epilepsy and paroxysmal dyskinesia (Du et al. 2005). The effect of this mutation is to shift the channel’s activation profile to the left, similar to the effect of inclusion of the STREX insert (Du et al. 2005; Yang et al. 2010). The STREX isoform also contributes to burst firing in hippocampal neurons (Bell et al. 2010). This suggests that the early predominance of the STREX isoform may increase seizure susceptibility. The β-4 subunit of the BK channel increases with postnatal development and deletion of the β-4 subunit results in seizures (Allen 2007; Brenner et al. 2005). Thus, lower expression of the β-4 subunit in early development may also contribute to seizure formation. Additionally, lower overall expression of the BK channel itself in interneurons may contribute to epilepsy as BK channels permit for high-frequency firing, a hallmark of interneuron activity which is necessary to regulate coordinated network activity in the hippocampus.

Understanding the implications of changes in BK expression and activity involves understanding these trends in the context of specific cellular subpopulations; the hippocampus comprises a diverse population of interneurons with varying roles in maintaining proper hippocampal excitability (for review see Pelkey et al. 2017). Changes in the excitability of a cell type could have differential effects on circuit excitability depending on the role of a that neuronal type in controlling the circuit excitability. Optogenetic and chemogenetic experiments have demonstrated that activating interneurons that directly inhibit pyramidal neurons (such as the parvalbumin and somatostatin expressing interneurons) can inhibit seizures, while activating cells that inhibit this first class of cells (such as VIP expressing interneurons) can potentiate excitatory transmission in the hippocampus (Cǎlin et al. 2018; Karnani et al. 2016). While some information about cell identity can be gleaned from their firing properties, that alone does not reliably differentiate cell populations (Wheeler et al. 2015). The use of firing properties alone is further confounded by the fact that action excitability changes rapidly in the early postnatal period. This work should be followed up by efforts to characterize trends in BK channel expression and activity in identified hippocampal neurons. Parvalbumin and somatostatin expressing interneurons are of particular interest since previous studies have demonstrated that activity in these interneurons may play a role in seizure activity (Cǎlin et al. 2018; Sessolo et al. 2015; Yekhlef et al. 2015). The immature properties and expression patterns of BK channels in these neurons may contribute to early-postnatal seizure susceptibility.

## Disclosures

The authors have no conflicts of interest of any kind to disclose.

## Funding

This work has been funded by the Marquette University Committee in Research and the Marquette University Department of Biological Sciences.

## Acknowledgements

The authors thank an undergraduate student, Alexis Monical, for her role in the initial data collection for this study. The authors also thank Dr. Deanna Arble for constructive feedback during the revision process.

## Notes

#### Summary of Updates

A couple of typographical errors were discovered on page 11. p> should have been p< in two places.

## References

Aika Y, Ren JQ, Kosaka K, Kosaka T. Quantitative analysis of GABA-like-immunoreactive and parvalbumin-containing neurons in the CA1 region of the rat hippocampus using a stereological method, the disector. Exp Brain Res 99: 267–276, 1994.

Allen. ALLEN Mouse Brain Atlas. Gene Expr, 2007. doi:10.1038/nature05453.

Banker GA, Cowan WM. Further observations on hippocampal neurons in dispersed cell culture. J Comp Neurol 187: 469–493, 1979.

Behrens R, Nolting A, Reimann F, Schwarz M, Waldscḧtz R, Pongs O. HKCNMB3 and hKCNMB4, cloning and characterization of two members of the large-conductance calcium-activated potassium channel β subunit family. FEBS Lett 474: 99–106, 2000.

Bell TJ, Miyashiro KY, Sul J-Y, Buckley PT, Lee MT, McCullough R, Jochems J, Kim J, Cantor CR, Parsons TD, Eberwine JH. Intron retention facilitates splice variant diversity in calcium-activated big potassium channel populations. Proc Natl Acad Sci 107: 21152–21157, 2010.

Brenner R, Chen QH, Vilaythong A, Toney GM, Noebels JL, Aldrich RW. BK channel beta4 subunit reduces dentate gyrus excitability and protects against temporal lobe seizures. Nat Neurosci 8: 1752–1759, 2005.

Cǎlin A, Jefferys JGR, Ilie AS, Akerman CJ, Stancu M, Zagrean A-M. Chemogenetic recruitment of specific interneurons suppresses seizure activity. Front Cell Neurosci 12, 2018.

Chen L, Tian L, MacDonald SHF, McClafferty H, Hammond MSL, Huibant JM, Ruth P, Knaus HG, Shipston MJ. Functionally diverse complement of large conductance calcium- and voltage-activated potassium channel (BK) α-subunits generated from a single site of splicing. J Biol Chem 280: 33599–33609, 2005.

Cline HT. Dendritic arbor development and synaptogenesis. Curr. Opin. Neurobiol. : 118–126, 2001.

Du W, Bautista JF, Yang H, Diez-Sampedro A, You SA, Wang L, Kotagal P, Lüders HO, Shi J, Cui J, Richerson GB, Wang QK. Calcium-sensitive potassium channelopathy in human epilepsy and paroxysmal movement disorder. Nat Genet 37: 733–738, 2005.

Dzhala VI, Staley KJ. Excitatory actions of endogenously released GABA contribute to initiation of ictal epileptiform activity in the developing hippocampus. J Neurosci 23: 1840–1846, 2003.

Galvez A, Gimenez-Gallego G, Reuben JP, Roy-Contancin L, Feigenbaum P, Kaczorowski GJ, Garcia ML. Purification and characterization of a unique, potent, peptidyl probe for the high conductance calcium-activated potassium channel from venom of the scorpion Buthus tamulus. J Biol Chem 265: 11083–11090, 1990.

Gao B-X, Ziskind-Conhaim L. Development of Ionic Currents Underlying Changes in Action Potential Waveforms in Rat Spinal Motoneurons. J Neurophysiol 80: 3047–3061, 2017.

Goldberg EM, Jeong HY, Kruglikov I, Tremblay R, Lazarenko RM, Rudy B. Rapid developmental maturation of neocortical FS cell intrinsic excitability. Cereb Cortex 21: 666–682, 2011.

Gómez-Di Cesare CM, Smith KL, Rice FL, Swann JW. Axonal remodeling during postnatal maturation of CA3 hippocampal pyramidal neurons. J Comp Neurol 384: 165–180, 1997.

Gu N, Vervaeke K, Storm JF. BK potassium channels facilitate high-frequency firing and cause early spike frequency adaptation in rat CA1 hippocampal pyramidal cells. J Physiol 580: 859–882, 2007.

Huttenlocher PR, Dabholkar AS. Regional differences in synaptogenesis in human cerebral cortex. J Comp Neurol 387: 167–178, 1997.

Jaffe DB, Wang B, Brenner R. Shaping of action potentials by type I and type II large-conductance Ca^2^+-activated K+ channels. Neuroscience 192: 205–218, 2011.

Kang J, Huguenard JR, Prince D a. Development of BK channels in neocortical pyramidal neurons. J Neurophysiol 76: 18–198, 1996.

Karnani MM, Jackson J, Ayzenshtat I, Sichani XH, Manoocheri K, Kim S, Yuste R. Opening holes in the blanket of inhibition: Localized lateral disinhibition by vip interneurons. J Neurosci 36: 3471–3480, 2016.

Kinnischtzke AK, Sewall AM, Berkepile JM, Fanselow EE. Postnatal maturation of somatostatin-expressing inhibitory cells in the somatosensory cortex of GIN mice. Front Neural Circuits 6, 2012.

Koschak A, Koch RO, Liu J, Kaczorowski GJ, Reinhart PH, Garcia ML, Knaus HG. [125I]iberiotoxin-D19Y/Y36F, the first selective, high specific activity radioligand for high-conductance calcium-activated potassium channels. Biochemistry 36: 1943–1952, 1997.

Kumar SS, Bacci A, Kharazia V, Huguenard JR. A developmental switch of AMPA receptor subunits in neocortical pyramidal neurons. J Neurosci 15: 3005–3015, 2002.

MacDonald SHF, Ruth P, Knaus HG, Shipston MJ. Increased large conductance calcium-activated potassium (BK) channel expression accompanied by STREX variant downregulation in the developing mouse CNS. BMC Dev Biol 6: 37, 2006.

McCormick DA, Prince DA. Postnatal development of electrophysiological properties of rat cerebral cortical pyramidal neurones. J Physiol 393: 743–762, 1987.

Meera P, Wallner M, Toro L. A neuronal β subunit (KCNMB4) makes the large conductance, voltage- and Ca2+-activated K+ channel resistant to charybdotoxin and iberiotoxin. Proc Natl Acad Sci U S A 97: 5562–5567, 2000.

Misonou H, Menegola M, Buchwalder L, Park EW, Meredith A, Rhodes KJ, Aldrich RW, Trimmer JS. Immunolocalization of the Ca2+-activated K+channel Slo1 in axons and nerve terminals of mammalian brain and cultured neurons. J Comp Neurol 496: 289–302, 2006.

Mynlieff M. Dissociation of postnatal hippocampal neurons for short term culture. J Neurosci Methods 73: 35–44, 1997.

Mynlieff M. Identification of different putative neuronal subtypes in cultures of the superior region of the hippocampus using electrophysiological parameters. Neuroscience 93: 479–486, 1999.

Pelkey KA, Chittajallu R, Craig MT, Tricoire L, Wester JC, McBain CJ. Hippocampal gabaergic inhibitory interneurons. Physiol Rev 97: 1619–1747, 2017.

Picken Bahrey HL, Moody WJ. Early development of voltage-gated ion currents and firing properties in neurons of the mouse cerebral cortex. J Neurophysiol 89: 1761–1763, 2003.

Rakhade SN, Jensen FE. Epileptogenesis in the immature brain: Emerging mechanisms. Nat. Rev. Neurol. 2009.

Sessolo M, Marcon I, Bovetti S, Losi G, Cammarota M, Ratto GM, Fellin T, Carmignoto G. Parvalbumin-positive inhibitory interneurons oppose propagation but favor generation of focal epileptiform activity. J Neurosci 35: 9544–9557, 2015.

Sheehan JJ, Benedetti BL, Barth AL. Anticonvulsant effects of the BK-channel antagonist paxilline. Epilepsia 50: 711–720, 2009.

Szczurowska E, Mareš P. NMDA and AMPA receptors: Development and status epilepticus. Physiol. Res. : S21–S38, 2013.

Wheeler DW, White CM, Rees CL, Komendantov AO, Hamilton DJ, Ascoli GA. Hippocampome.org: A knowledge base of neuron types in the rodent hippocampus. Elife 4: e09960, 2015.

Xie J, McCobb DP. Control of alternative splicing of potassium channels by stress hormones. Science (80-) 280: 443–446, 1998.

Yang J, Krishnamoorthy G, Saxena A, Zhang G, Shi J, Yang H, Delaloye K, Sept D, Cui J. An Epilepsy/Dyskinesia-Associated Mutation Enhances BK Channel Activation by Potentiating Ca2+ Sensing. Neuron 66: 871–883, 2010.

Yekhlef L, Breschi GL, Lagostena L, Russo G, Taverna S. Selective activation of parvalbumin or somatostatin-expressing interneurons triggers epileptic seizurelike activity in mouse medial entorhinal cortex. J Neurophysiol 113: 1616–1630, 2015.

Zhou X-B, Wulfsen I, Utku E, Sausbier U, Sausbier M, Wieland T, Ruth P, Korth M. Dual role of protein kinase C on BK channel regulation. Proc Natl Acad Sci 107: 8005–8010, 2010.

Zhou X, Wulfsen I, Korth M, McClafferty H, Lukowski R, Shipston MJ, Ruth P, Dobrev D, Wieland T. Palmitoylation and membrane association of the stress axis regulated insert (STREX) controls BK channel regulation by protein kinase C. J Biol Chem 287: 32161–32171, 2012.

